# Border-ownership-dependent tilt aftereffect for shape defined by binocular disparity and motion parallax

**DOI:** 10.1101/472076

**Authors:** Reuben Rideaux, William J Harrison

## Abstract

Discerning objects from their surrounds (i.e., figure-ground segmentation) in a way that guides adaptive behaviours is a fundamental task of the brain. Neurophysiological work has revealed a class of cells in the macaque visual cortex that may be ideally suited to support this neural computation: border-ownership cells (Zhou, Friedman, & von der Heydt, 2000). These orientation-tuned cells appear to respond conditionally to the borders of objects. A behavioural correlate supporting the existence of these cells in humans was demonstrated using two-dimensional luminance defined objects (von der Heydt, Macuda, & Qiu, 2005). However, objects in our natural visual environments are often signalled by complex cues, such as motion and depth order. Thus, for border-ownership systems to effectively support figure-ground segmentation and object depth ordering, they must have access to information from multiple depth cues with strict depth order selectivity. Here we measure in humans (of both sexes) border-ownership-dependent tilt aftereffects after adapting to figures defined by either motion parallax or binocular disparity. We find that both depth cues produce a tilt aftereffect that is selective for figure-ground depth order. Further, we find the effects of adaptation are transferable between cues, suggesting that these systems may combine depth cues to reduce uncertainty (Bülthoff & Mallot, 1988). These results suggest that border-ownership mechanisms have strict depth order selectivity and access to multiple depth cues that are jointly encoded, providing compelling psychophysical support for their role in figure-ground segmentation in natural visual environments.

**SIGNIFICANCE STATEMENT:** Segmenting a visual object from its surrounds is a critical function that may be supported by “border-ownership” neural systems that conditionally respond to object borders. Psychophysical work indicates these systems are sensitive to objects defined by luminance contrast. To effectively support figure-ground segmentation, however, neural systems supporting border-ownership must have access to information from multiple depth cues and depth order selectivity. We measured border-ownership-dependent tilt aftereffects to figures defined by either motion parallax or binocular disparity and found aftereffects for both depth cues. These effects were transferable between cues, but selective for figure-ground depth order. Our results suggest that the neural systems supporting figure-ground segmentation have strict depth order selectivity and access to multiple depth cues that are jointly encoded.

## INTRODUCTION

Our natural visual environments are complex and often cluttered with objects. In order to interact with our surrounds appropriately, therefore, objects must be segmented from other objects and their backgrounds, and their position in depth must be inferred from often fragmented and ambiguous cues such as binocular disparity, motion parallax, and texture. Achieving such so-called “figure-ground segmentation” with the speed and automaticity necessary to effectively function within the environment is non-trivial; understanding how the brain accomplishes this remains of fundamental importance in neuroscience.

Neurophysiological work has revealed a class of cells in the macaque primary visual cortex that has been implicated in figure-ground segmentation. These cells, known as border-ownership cells (Zhou et al., 2000), are tuned to oriented edges, like many neurons in the primary visual cortex (Hubel & Wiesel, 1959). However, unlike most orientation-tuned cells, the activity of border-ownership cells is contingent on whether or not the edge belongs to the border of an object (Zhou et al., 2000). For instance, the same light-dark edge presented within the receptive field of a neuron could produce a larger increase in firing rate if the light region was part of a distinct ‘figure’ positioned on a dark ‘background’ than vice versa (**Fig. 1a**). Critically, this contingency is active even when the object extends far beyond the classic receptive field of the cell, suggesting that border-ownership cells are connected through a network that can identify borders that are common to an object. This distinctive characteristic is ideally suited for binding the borders of an object in order to segment it from the background and facilitate other figure-ground mechanisms to retrieve those borders as a whole.

**Figure 1.**
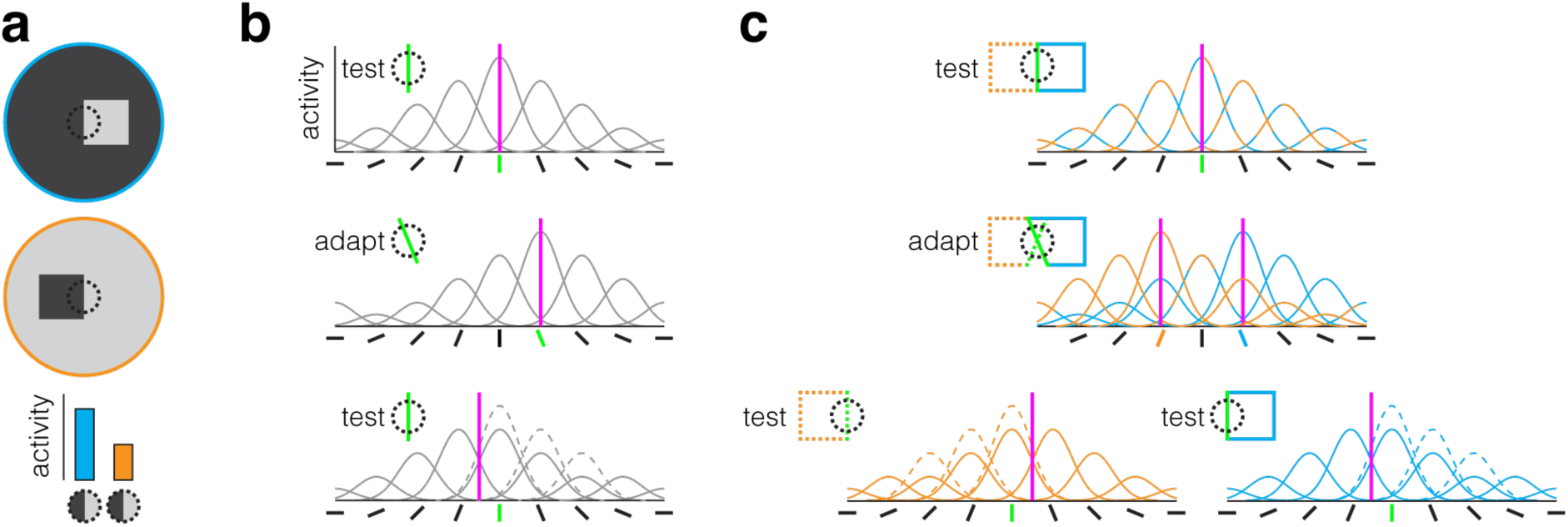
Illustrations of border-ownership neurons and putative explanations of tilt-aftereffects. (**a**) The same dark-light edge within the (dotted circle) receptive field of certain orientation-tuned neurons will evoke different levels of activity depending on whether (cyan) the light region is part of a “figure” and the dark region part of the “ground” or (orange) vice versa. (**b**) The (magenta line) perceived orientation of a (green) vertical bar is based on (top) the population response of multiple orientation tuned cells. Prolonged exposure to a bar tilted clockwise from vertical reduces (middle) the responsiveness of cells tuned to this orientation (i.e., adaptation). Following adaptation, viewing a vertical bar will now produce (bottom) a population response that is biased away from the adapted orientation, i.e., a tilt-aftereffect, due to (dashed lines) the attenuated response of the cells. (**c**) By extending the vertical bar such that it becomes the border of a figure, on (dotted orange) the left or on (cyan) the right, (top) the bar will now evoke activity from separate populations of border-ownership cells. Prolonged alternating viewing of borders tilted clockwise and anti-clockwise from vertical that belong to the left and right figures, respectively, will produce (middle) border-ownership-dependent adaptation in separate populations of cells with receptive fields in the same retinotopic location. Now, when a vertical border is viewed it will produce (bottom) a population response that is biased away from the adapted tilt orientation of the figure it belongs.

A behavioural correlate of the contingency that characterizes border-ownership cells can be demonstrated psychophysically in humans using a modification of the classic tilt-aftereffect paradigm (**Fig. 1b**). This was elegantly demonstrated by von der Heydt, Macuda, and Qiu (2005), who presented two luminance-defined adaptor figures alternating in time. One object was presented to the left of centre, while the other object was presented to the right of centre. Importantly, the inner border of these objects intersected in space, but were of opposite tilt. When a solitary test line was subsequently presented at the intersection, no tilt-aftereffect was observed, presumably because the effects of the adapting edges were balanced. In contrast, if the test line was extended to form a figure to the left or the right of fixation, a tilt-aftereffect was observed that was repulsed away from the border of the adaptor on the side to which the test line extended (**Fig. 1c**).

Objects in our natural environment are seldom defined by a simple luminance discontinuity; their borders are often signalled by multiple depth cues such as motion parallax and binocular disparity. Thus, for the neural systems of border-ownership to effectively support figure-ground segmentation, they must have access to information from multiple depth cues. Further, in order to establish the depth order of borders these systems must have depth order selectivity. Note, while border-ownership (i.e., the property of an edge which denotes whether it borders an object or not) may also indicate depth order (i.e., the specific depth configuration of an edge), this is not always the case. For instance, cells can be selective for one but not the other, and in the absence of disambiguating cues, the depth order of borders can be ambiguous, e.g., Rubin’s face-vase illusion. Finally, depth cues are noisy and often unreliable; in order to yield accurate inferences, psychophysical evidence suggests that for some tasks the brain combines information from multiple depth cues (Ernst & Bülthoff, 2004). Thus, if depth cues are used by the neural systems supporting border-ownership, determining whether these cues are jointly encoded will reveal if similar mechanisms for uncertainty reduction may be utilized in figure-ground segmentation.

To test whether the neural systems supporting border-ownership use depth cues, and whether they are selectivity tuned to a particular depth order, we measured border-ownership-dependent tilt-aftereffects following adaptation to figures defined by two primary depth cues: 1) motion parallax and 2) binocular disparity. We found that both depth cues produce a tilt-aftereffect which is selective for depth order. Moreover, we found that the effect of adaptation is transferable between cues. These results indicate that the neural systems supporting perceptual figure-ground segmentation have strict depth order selectivity and access to multiple depth cues that are jointly encoded.

## METHODS

### Participants

Observers were recruited from the University of Cambridge and had normal or corrected-to-normal vision and were screened for stereo deficits using a fine discrimination task (just-noticeable difference < .5 arcmin). Eight right-handed human adults participated in each of the two experiments (Experiment 1: 4 male, 26.3±5.6 years; Experiment 2: 3 male, 26.7±4.9 years). Five participants participated in both experiments, including one author (RR). With the exception of (RR), all participants were naïve to the aims of the experiment. Experiments were approved by the University of Cambridge ethics committee; all observers provided written informed consent.

### Apparatus and stimuli

Stimuli were generated in MATLAB (The MathWorks, Inc., Matick, MA) using Psychophysics Toolbox and Eyelink Toolbox extensions (Brainard, 1997; Cornelissen, Peters, & Palmer, 2002; Pelli, 1997; see http://psychtoolbox.org/). Binocular presentation was achieved using a pair of Samsung 2233RZ LCD monitors (120Hz, 1680×1050) viewed through front-silvered mirrors in a Wheatstone stereoscope configuration. The viewing distance was 50 cm and the participants’ head position were stabilized using an eye mask, headrest and chin rest. Eye movement was recorded binocularly at 1 kHz using an EyeLink 1000 (SR Research Ltd., Ontario, Canada).

Stimuli and design were motivated by those used by (von der Heydt et al., 2005). Unlike this previous study that used solid lines on a blank background, however, our stimuli consisted of pixel noise in which shapes were distinguished by either (i) binocular disparity or (ii) motion parallax, appearing as (near) a “figure” or (far) a “window” in front of a background surface (**Fig. 2a**). Binocular disparity-defined shapes could be either near or far (±2.5 arcmin) relative to the background, with the closer plane always at zero degrees offset (**Fig. 2b**). Pixel (white, 198 cd/m^2^; black, 5 cd/m^2^) luminance was randomly reassigned at 10 Hz. Similarly, motion parallax-defined shapes were made to appear near or far by moving the pixels on either the inside or the outside of the trapezoidal region horizontally according to a sinusoidal function (max speed 40/s, with an initial phase 0 or pi, counterbalanced across presentations; **Fig. 2c**). To reduce non-target cues to orientation that could be used as a reference, the stimulus region was circular (radius 4°) with a Gaussian smoothed edge (sd 8 arcmin). Trapezoidal regions (height 3°, centre width 3°) were positioned on the left or the right side of the centre of the screen with the tilted edge (henceforth referred to as the flank) centred on the midline. A blue dot (radius 6 arcmin) was positioned to the left or right of centre (30 armin) to stabilize fixation. During blank periods the stimulus region was mid-grey (99 cd/m^2^); the remainder of the screen was always black (5 cd/m^2^).

**Figure 2.**
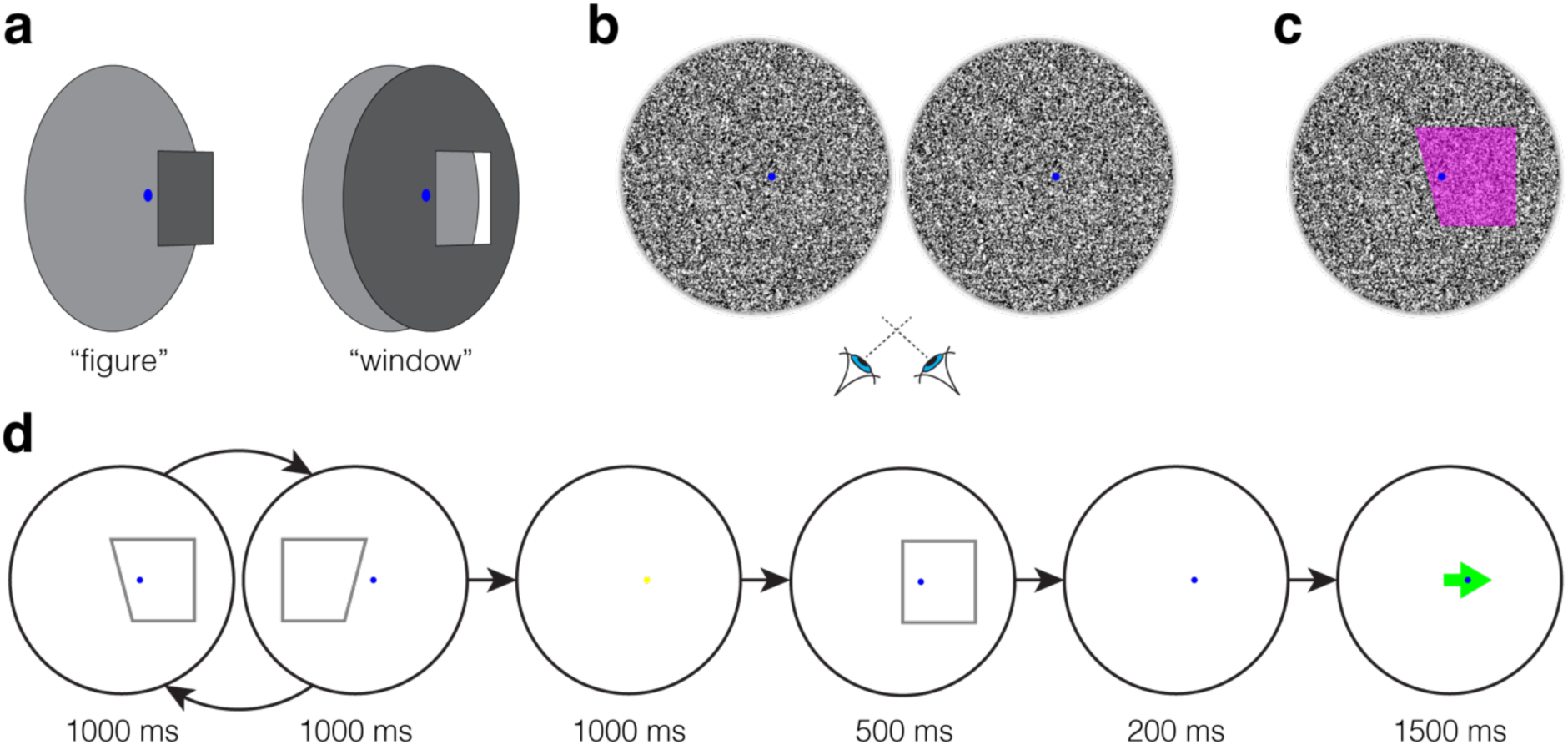
Examples of experimental stimuli and design. (**a**) Illustration of the two (figure/window) depth structures used in the experiment. (**b**) Stereogram of the (figure) adaptor stimulus in the binocular disparity condition designed for cross-eyed fusion. (**c**) One frame of the adaptor stimulus in the motion parallax condition; the figure region is indicated in magenta. Pixels either (window) inside or (figure) outside the trapezoidal region were moved horizontally according to a sinusoidal function.(**d**) Schematic of the design; alternating presentation of adaptation figures on the left and right (adaptation sequence) followed by a test figure and an arrow pointing either left or right, each separated by blank intervals. The observers’ task was to indicate whether the orientation of the test figure flank was consistent with the direction of the proceeding arrow.

### Procedure

Runs consisted of an initial adaptation sequence (32 pairs of trapezoids; 76.8 s), followed by test trials that were separated by top-up adaptation sequences (three pairs of trapezoids; 7.2 s; **Fig. 2d**). During the adaptation sequences, observers viewed trapezoidal figures (left and right flank tilt either [−15, 15]° or [15, −15]° in separate blocks) alternating between left- and right-side presentation (duration 1 s, ISI .2 s). The depth cue (binocular disparity or motion parallax) that defined the adaptor and test figures was always the same within a run, but the depth order could be either congruent (e.g., figure with figure) or incongruent (e.g., figure with window). A method of constant stimuli was used with trials consisting of a test figure (flank tilt ±[8, 4.8, 1.6]°) presented pseudo-randomly, either on the left or the right of the screen centre for a duration of 0.5 s. Following the test stimulus, there was a 0.2 s blank period followed by presentation of a green arrow centred on fixation, pointing left or right (1.5 s). To mitigate fixation disruption, the fixation dot always occluded the arrow. The arrow was presented at the same depth plane as the fixation dot, and its direction was selected at random.

Observers’ task was to press the “space” key when the test flank’s tilt direction (anticlockwise/left or clockwise/right) appeared to be consistent with the direction of the following arrow. This method of response was used instead of separate left/right responses to avoid systematic response bias, such as observers pressing left or right more frequently under high uncertainty. A 1 s blank interval separated adaptation and test periods to avoid a potential bias of the afterimage of the last-presented adaptation figure. For the same reason, the starting trapezoid of the adaptation sequence was alternated so that left-/right-side trapezoids were presented as the last figure on half the trials. During the blank interval preceding each trial, the fixation dot was changed from blue to yellow to prepare observers for the upcoming trial. Prior to adaptation runs, baseline test runs were performed in which subjects were presented with the same test stimuli without adaptation.

In Experiment 1, we used a 2×2×2 factorial design: depth cue (motion parallax/binocular disparity) × depth (figure/window) × depth congruence (congruent/incongruent). Adaptation and baseline runs consisted of 48 and 72 trials, respectively. For each condition, observers underwent two baseline and two adaptation runs; one for each of the left-/right-positioned fixation dot (randomized order). In each session, observers completed two conditions, for a total duration of approximately 60 min. To avoid carry-over effects, sessions were completed on separate days and the adaptation orientation was reversed between conditions. Congruent runs were completed prior to incongruent runs and the depth cue condition was held constant within sessions. The order of depth sign and depth cue conditions was counterbalanced across participants.

In Experiment 2, we tested whether adaptation to one depth cue influenced objects defined by the other depth cue. We therefore ran two conditions where the depth portrayed a (near) figure, but the cue used to define the adaptor and test stimuli were incongruent. For example, an observer would adapt to motion-defined objects, and were then tested with disparity-defined objects, or vice versa.

### Task response data analysis

For each condition, we concatenated data from the two runs and fit it with a psychometric function using the MATLAB toolbox Psignifit (Frund, Haenel, & Wichmann, 2011; see http://psignifit.sourceforge.net/) to establish separate threshold, i.e., point of subjective equality (PSE), and slope values for responses corresponding to test figures presented on the left and right. To determine the bias on perceived orientation that resulted from adaptation, we calculated the difference between baseline and adaptation threshold values. To match the bias between runs with [−15, 15]° tilt adaptors across observers, we reversed the sign of bias in the [15, −15]° tilt runs. For the same reason, we reversed the sign of the bias for figures presented on the left prior to averaging the two, yielding a single measure of bias for each condition. Thus, average bias was normalized to the −15° tilt adaptor condition.

### Eye tracking data analysis

Prior to analysis, eye movement data were screened to remove blinks and noisy and/or spurious recordings. To test for differences in eye position between experimental conditions, we calculated the mean and standard deviation of observers’ vertical and horizontal binocular eye position, relative to fixation, for version and vergence eye movements during adaptor and test presentation. To match eye position coordinates between runs with left- and right-side fixation, we reversed the sign of horizontal eye movements in runs where the fixation dot was presented on the left side.

## RESULTS

### Selectivity for multiple depth cues

Neurophysiological work has revealed orientation-tuned cells in the primary visual cortex of macaque that conditionally respond to the borders of figures. Psychophysical work with humans has demonstrated a behavioural correlate of this cellular characteristic: border-ownership-dependent tilt aftereffects (von der Heydt et al., 2005). For a neural system to effectively support figure-ground segmentation in a three-dimensional environment, there are at least two crucial characteristics required: 1) sensitivity to multiple depth cues and 2) depth order preference.

Here we tested whether adaptation to figures defined by one of two primary depth cues (motion parallax or binocular disparity) produce a border-ownership-dependent tilt-aftereffect. We further investigated whether any adaptation effect is transferable between depth order configurations (e.g., adapt to a near object, and test with a far object). We found that perceptual bias following adaptation was significantly larger when the adaptor and test stimulus depth were congruent than incongruent (repeated-measures ANOVA; F_1,49_=37.6, P=1.5e^-7^, *d*=4.3.; **Fig. 3a & b**). Indeed, we found a significant adaptation effect for all congruent conditions, but failed to find an effect for any incongruent condition (**Fig. 3b**). We also found no difference in the strength of the effect for different cues (F_1,49_=0.4, P=.521, *d*=0.4) or depths (F_1,49_=4.0, P=.087, *d*=1.4). No interactions were significant (all ps>.05). These results indicate that border-ownership cells are sensitive to both motion parallax and binocular disparity, and that they show depth order preference. A possible concern is that we failed to detect a bias in the incongruent conditions because bias estimates were less precise. Such a problem could arise because of, for example, reduced anticipation of the test stimulus due to the change in depth relative to the cue stimulus. However, we found no evidence for a difference in the precision of observers’ estimates between congruent and incongruent conditions (F_1,49_=0.04, P=.849, *d*=0.14; **Fig. 3c**). By contrast, we found that observers’ judgements were more precise for stimuli defined by motion parallax than binocular disparity (F_1,49_=15.6, P=.006, *d*=2.8). No differences in the precision of estimates were observed between depth conditions (F_1,49_=4.2, P=.079, *d*=1.4).

**Figure 3.**
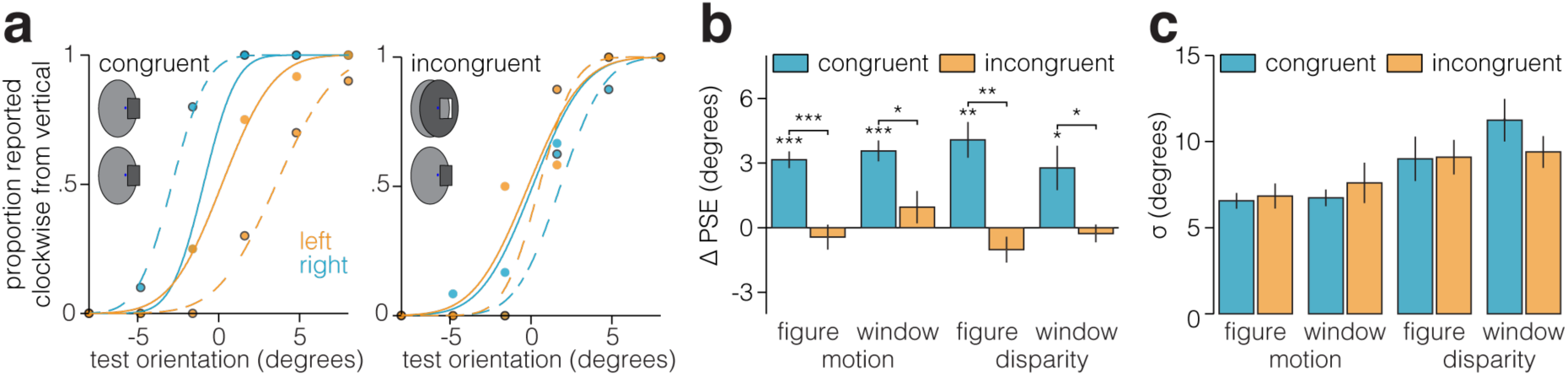
Psychophysical measurements of orientation judgements following adaptation. Following adaptation to oriented stimuli defined by either motion parallax or binocular disparity, observers judged the orientation of the edge of a test stimulus that was either the same (congruent) or different (incongruent) depth order. (**a**) A representative example of the psychometric functions fit to one (naïve) participants’ data in the motion-figure congruent and incongruent conditions. Solid lines, and dots without outlines, indicate baseline data, dashed lines and dots with black outlines indicate adaptation data. Orange and cyan dots and lines indicate data from test figures presented on the left and right sides, respectively. (**b**) Relative to a pre-adaptation baseline, observers’ point of subjective equality (PSE) was biased away from the orientation of the adaptor when the test stimulus was congruent (paired t-test; motion-near, t_7_=8.8, P=4.9e^-5^, *d*=6.2; motion-far, t_7_=7.9, P=9.5e^-5^, *d*=5.6; disparity-near, t_7_=5.3, P=.001, *d*=3.7; disparity-far, t_7_=2.8, P=.023, *d*=2.0), but not incongruent (paired t-test; motion-near, t_7_=-0.8, P=.446, *d*=0.6; motion-far, t_7_=1.3, P=.216, *d*=0.9; disparity-near, t_7_=-1.8, P=.120, *d*=1.2; disparity-far, t_7_=-0.7, P=.508, *d*=0.5), with the adaptor. Further, the PSE in congruent conditions was significantly more biased than in corresponding incongruent conditions (paired t-test; motion-near, t_7_=2.7, P=.029, *d*=1.9; motion-far, t_7_=6.9, P=2.3e^-4^, *d*=4.9; disparity-near, t_7_=4.5, P=.003, *d*=3.2; disparity-far, t_7_=2.8, P=.028, *d*=2.0). The difference in bias between congruent and incongruent conditions cannot be explained by differences in the precision of judgements, (**c**) we found no differences in precision (inverse of sigma) between corresponding in/congruent conditions (paired t-test; motion-near, t_7_=0.7, P=.513, *d*=0.5; motion-far, t_7_=0.8, P=.420, *d*=0.6; disparity-near, t_7_=0.1, P=.956, *d*=0.1; disparity-far, t_7_=1.6, P=.152, *d*=1.3). Note that labels along the abscissa refer to the depth of the test stimulus.

The border-ownership-dependent tilt-aftereffect is retinotopically localized to a relatively small region (~2°)(von der Heydt et al., 2005); thus, another possible concern is that we failed to detect a tilt-aftereffect in the incongruent conditions because observers’ gaze position moved more between adaptor and test stimulus presentation than in congruent conditions. However, we found no evidence for a larger change in binocular eye position between adaptor and test periods for either horizontal/vertical vergence (RM ANOVA; horizontal, F_1,49_=2.8, P=.159, *d*=1.2; vertical, F_1,49_=0.7, P=.433, *d*=0.6; **Fig. 4a**) or version eye movements (horizontal, F_1,49_=0.4, P=.541, *d*=0.4; vertical, F_1,49_=0.3, P=.593, *d*=0.4; **Fig. 4b**).

**Figure 4.**
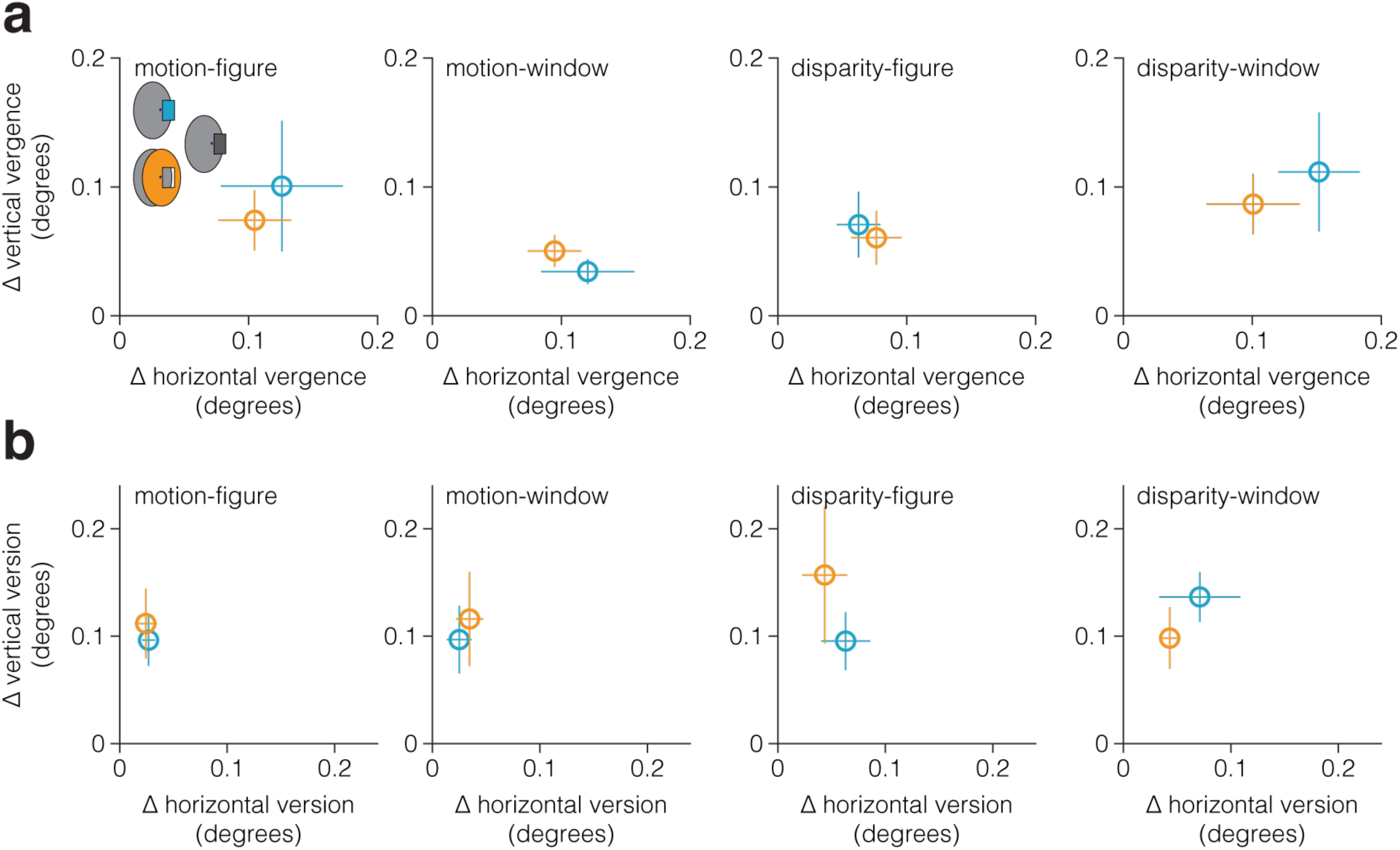
Change in binocular eye gaze position between adaptor and test stimulus presentation. To test whether our failure to detect transference of adaptation between (relatively) near and far figure borders was due to larger changes in eye gaze position between test and adaptor stimulus presentation in the incongruent condition, we compared the absolute difference in the average horizontal/vertical (**a**) vergence and (**b**) version eye movements between (cyan) congruent and (orange) incongruent conditions. We found no evidence for a larger change in binocular eye gaze position for either vergence or version eye movements. Note, the caption at the top of each plot indicates the test stimulus condition.

### Joint encoding of depth cues

The results above suggest that the neural systems supporting border-ownership are tuned to motion parallax and binocular disparity depth cues. There are two ways in which these cues may be encoded by these systems: 1) depth cues are encoded in separate neural populations, or 2) neurons supporting border-ownership jointly encode depth cues. Consistent with the joint encoding hypothesis, previous neurophysiological work indicates that some border-ownership cells combine binocular disparity and Gestalt cues (Qiu & von der Heydt, 2005). To determine whether the neural systems supporting border-ownership encode (motion parallax and binocular disparity) depth cues separately or jointly, we tested whether the effect of adaptation to a stimulus defined by one cue was transferred to a stimulus defined by the other. In line with the hypothesis that cues are jointly encoded, we found a significant tilt-aftereffect for stimuli defined by motion parallax, following adaptation to stimuli defined by binocular disparity (paired t-test, t_7_=6.3, P=3.9e^-4^, *d*=4.4), and vice versa (t_7_=8.0, P=8.8e^-5^, *d*=5.7; **Fig. 5a**). Consistent with our previous results, we again found that judgements for stimuli defined by motion parallax were more precise (t_7_=4.1, P=.004, *d*=2.9; **Fig. 5b**).

**Figure 5.**
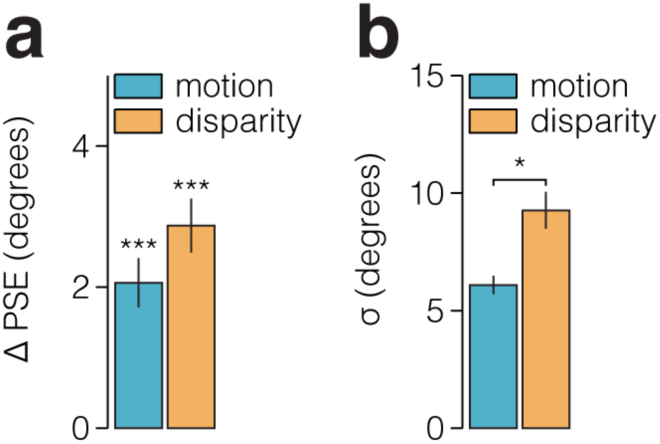
Transfer of adaptation effects between stimuli defined by motion parallax and binocular disparity. Following adaptation to oriented figure defined by either motion parallax or binocular disparity, observers judged the orientation of a test figure that was defined by the other cue.(**a**) Relative to a pre-adaptation baseline, observers’ point of subjective equality (PSE) was biased away from the orientation of the adaptor for both stimuli. Note that the shade of the bars indicates the depth cue used to define the test stimulus, whereas the opposite cue was the adaptor stimulus. (**b**) Consistent with the results from the previous experiment, we also found that judgements made for stimuli defined by motion were more precise than for disparity.

## DISCUSSION

Psychophysical work with human observers has shown that multiple tilt aftereffects can be induced at the same retinotopic location by adapting to the orientation of overlapping borders of different figures (von der Heydt et al., 2005). This finding supports evidence from neurophysiological work with macaque revealing orientation-tuned border-ownership cells in the primary visual cortex that conditionally respond to the figure borders (Zhou et al., 2000). These findings were obtained using luminance-defined objects viewed on two-dimensional displays; however, objects in our natural environment are seldom defined in this manner, they are often signalled by multiple depth cues such as motion parallax and binocular disparity. To efficiently segment the visual environment the neural systems supporting border-ownership would be expected to have access to depth cues and to show strong depth order tuning. To test this hypothesis, here we measured border-ownership-dependent tilt-aftereffects for stimuli defined by two primary depth cues, motion parallax and binocular disparity. We found tilt-aftereffects for both cues, which were *not* transferable between depth order configurations (e.g., adapting to a “figure” edge did not influence perception of a “window” edge), but were transferable between cues (e.g., adapting to motion parallax influenced perception of a subsequent object defined by binocular disparity).

The finding that a border-ownership-dependent tilt aftereffect can be induced using stimuli defined by motion parallax or binocular disparity shows that the neural systems supporting border-ownership have access to information from multiple depth cues. These psychophysical results provide converging evidence consistent with previous work showing that binocular disparity and Gestalt cues are both encoded by border-ownership cells (Qiu & von der Heydt, 2005) and may indicate that border-ownership cells are sensitive to a range of depth cues. The finding that tilt aftereffects were non-transferable between depth configurations shows that the neural systems supporting border-ownership have strict depth order preference; an essential characteristic for establishing the depth order of objects within a scene.

Given most objects that are the focus of the visual system are figure-like rather than window-like, one may expect aftereffects to be stronger for the borders of figures than windows. However, we found no evidence to support this hypothesis; there was no difference between the strength of aftereffects for figures and windows defined by either motion parallax or binocular disparity.

If the behavioural effects observed here are a result of adapting the responses of a population of border-ownership cells in the human brain, it is interesting to consider how our results align with neurophysiological evidence of these cells in macaque. For instance, Qiu and von der Heydt (2005) found that only just over half of border-ownership cells tuned to binocular disparity were also depth order selective. This seems (at least partially) at odds with the strict depth order preference of the tilt aftereffects. That is, if a considerable proportion (40%) of border-ownership cells are not selective for depth order, it may have been reasonable to have expected to find transference of adaptation between depth order conditions; yet, we found no evidence for this. If the behavioural effects observed here were produced by adapting border-ownership cells, this may suggest that depth order selectivity is more prevalent in these cells than previously estimated. However, the proportion of depth order selective border-ownership cells in the macaque primary visual cortex may not be representative of those in the human’s, and the relationship between the proportion of these cells and their behavioural consequences may not be linear. Further, in the present experiments, observers likely attended to the figures, whereas in the macaque experiments the subjects’ attention was likely focused on the fixation task (Qiu & von der Heydt, 2005); thus, differences across species may be due to differences in attentional allocation across tasks (Fang, Boyaci, & Kersten, 2009; Qiu, Sugihara, & von der Heydt, 2007). This cross-species incongruence underscores the importance of psychophysical evidence in establishing the perceptual influence of stimuli as processed by an entire neural system.

The finding that adaptation effects could be transferred between stimuli defined by either motion parallax or binocular disparity indicates that the neural systems supporting border-ownership jointly encode these depth cues. Border-ownership mechanisms serve to bind contour features to larger entities and have a high degree of orientation and spatial position specificity. In contrast, the higher level figure-ground mechanisms first demonstrated by Lamme (1995) that support the response modulation observed between figure and ground regions are, among other differences summarized by von der Heydt (2015), less spatially precise and occur independently of the tuning properties of neurons (Zipser, Lamme, & Schiller, 1996). The present findings suggest that the responses of a subpopulation of neurons tuned to a particular orientation were reduced contingent on specific left/right and near/far positional characteristics of the figure. This specificity is characteristic of border-ownership mechanisms, indicating that integration of motion parallax and binocular disparity signals is supported by early border-ownership neural systems, rather than later figure-ground processes. The current findings cannot distinguish whether integration is performed by border-ownership cells or by higher level “groupings cells” (Craft et al., 2007). However, the finding that binocular disparity and Gestalt cues are integrated by border-ownership cells (Qiu & von der Heydt, 2005) implicates this earlier stage as a likely candidate.

Functional MRI work with humans indicates that motion parallax and depth cues are combined in area V3B/KO (Ban, Preston, Meeson, & Welchman, 2012). By contrast, neurophysiological work has identified border-ownership cells in area V2 of the macaque cortex that jointly encode binocular disparity and Gestalt cues (Qiu & von der Heydt, 2005). Our results may therefore indicate that motion parallax and binocular disparity signals are combined prior to V3B/KO. For instance, the neural systems that combine the signals evoked by these cues may be present as early as V2 and culminate in higher cortical areas such as V3B/KO where they are sufficiently prevalent to identify with fMRI. Alternatively, these results may indicate that border-ownership cells, which combine motion parallax and binocular disparity, are present in V3B/KO. Given that neurophysiological work has exclusively found evidence for border-ownership cells in ventral regions (V2 & V4), the presence of these cells in dorsal area V3B/KO would suggest they play a more ubiquitous role visual processing than previously thought.

Several models have been proposed to capture the mechanism by which border-ownership systems perform figure-ground segmentation, which can be broadly categorized as either feedforward (Sakai & Nishimura, 2006; Supèr, Romeo, & Keil, 2010), lateral (Zhaoping, 2005), or feedback models (Craft, Schutze, Niebur, & von der Heydt, 2007; Jehee, Lamme, & Roelfsema, 2007); see Williford and Heydt (2013) for a review. Our data provide new challenges for these models, as they will need to incorporate the capacity of border-ownership systems to jointly encode multiple depth cues with a high prevalence of depth order selectivity. Similarly, multiple models exist for describing the way in which the brain combines cues (Ohshiro, Angelaki, & Deangelis, 2011; Rideaux & Welchman, 2018); given that border-ownership mechanisms appear to combine depth cues, future work may also investigate how border-ownership cells combine cues and whether it can be described by an existing model, e.g. maximum likelihood estimation (Bülthoff & Mallot, 1988) or Bayesian (Knill & Pouget, 2004) frameworks.

Previous work demonstrated a behavioural correlate of border-ownership using two-dimensional luminance defined objects (von der Heydt et al., 2005). The current study exploits this correlate to show that the neural systems supporting border-ownership in humans jointly encode multiple depth cues with strong depth order selectivity. These results provide compelling evidence for the capacity of border-ownership mechanisms to support figure-ground segmentation in natural visual environments. Further, these data provide an important future question for neurophysiological work to pursue; that is, the nature and locus of border-ownership cells that combine motion parallax and binocular disparity signals.

## Acknowledgements

This work was supported by the Leverhulme Trust (ECF-2017-573 to RR) and the Wellcome Trust (095183/Z/10/Z to RR as a postdoctoral fellow of Andrew E Welchman), and by a National Health and Medical Research Council of Australia (CJ Martin Fellowship APP1091257 to WJH).

## REFERENCES

Ban, H., Preston, T. J., Meeson, A., & Welchman, A. E. (2012). The integration of motion and disparity cues to depth in dorsal visual cortex. Nature Neuroscience, 15(4), 636–643. https://doi.org/10.1038/nn.3046

Brainard, D. H. (1997). The Psychophysics Toolbox. Spatial Vision, 10(4), 433–436. https://doi.org/10.1163/156856897X00357

Bülthoff, H. H., & Mallot, H. a. (1988). Integration of depth modules: stereo and shading. Journal of the Optical Society of America. A, Optics and Image Science, 5(10), 1749-1758. https://doi.org/10.1364/JOSAA.5.001749

Cornelissen, F. W., Peters, E. M., & Palmer, J. (2002). The Eyelink Toolbox: eye tracking with MATLAB and the Psychophysics Toolbox. Behavior Research Methods, Instruments & Computers, 34(4), 613–617.

Craft, E., Schutze, H., Niebur, E., & von der Heydt, R. (2007). A Neural Model of Figure-Ground Organization. Journal of Neurophysiology, 97(6), 4310–4326. https://doi.org/10.1152/jn.00203.2007

Ernst, M. O., & Bülthoff, H. H. (2004). Merging the senses into a robust percept. Trends in Cognitive Sciences, 8(4), 162–169. https://doi.org/10.1016/j.tics.2004.02.002

Fang, F., Boyaci, H., & Kersten, D. (2009). Border Ownership Selectivity in Human Early Visual Cortex and its Modulation by Attention. Journal of Neuroscience, 29(2), 460–465. https://doi.org/10.1523/JNEUROSCI.4628-08.2009

Frund, I., Haenel, N. V., & Wichmann, F. A. (2011). Inference for psychometric functions in the presence of nonstationary behavior. Journal of Vision, 11(6), 16–16. https://doi.org/10.1167/11.6.16

Hubel, D. H., & Wiesel, T. N. (1959). Receptive fields of single neurones in the cat’s striate cortex. The Journal of Physiology, 148(3), 574-591. https://doi.org/10.1113/jphysiol.1959.sp006308

Jehee, J. F. M., Lamme, V. A. F., & Roelfsema, P. R. (2007). Boundary assignment in a recurrent network architecture. Vision Research, 47(9), 1153–1165. https://doi.org/10.1016/j.visres.2006.12.018

Knill, D. C., & Pouget, A. (2004). The Bayesian brain: The role of uncertainty in neural coding and computation. Trends in Neurosciences, 27(12), 712–719. https://doi.org/10.1016/j.tins.2004.10.007

Lamme, V. A. (1995). The neurophysiology of figure-ground segregation in primary visual cortex. The Journal of Neuroscience?: The Official Journal of the Society for Neuroscience, 15(2), 1605–1615. https://doi.org/10.1523/JNEUROSCI.15-02-01605.1995

Ohshiro, T., Angelaki, D. E., & Deangelis, G. C. (2011). A normalization model of multisensory integration. Nature Neuroscience, 14(6), 775–782. https://doi.org/10.1038/nn.2815

Pelli, D. G. (1997). The VideoToolbox software for visual psychophysics: Transforming numbers into movies. Spatial Vision. https://doi.org/10.1163/156856897X00366

Qiu, F. T., Sugihara, T., & von der Heydt, R. (2007). Figure-ground mechanisms provide structure for selective attention. Nature Neuroscience, 10(11), 1492–1499. https://doi.org/10.1038/nn1989

Qiu, F. T., & von der Heydt, R. (2005). Figure and ground in the visual cortex: V2 combines stereoscopic cues with Gestalt rules. Neuron, 47(1), 155–156. https://doi.org/10.1016/j.neuron.2005.05.028

Rideaux, R., & Welchman, A. E. (2018). Proscription supports robust perceptual integration by suppression in human visual cortex. Nature Communications, 9(1), 1502. https://doi.org/10.1038/s41467-018-03400-y

Sakai, K., & Nishimura, H. (2006). Surrounding Suppression and Facilitation in the Determination of Border Ownership. Journal of Cognitive Neuroscience, 18(4), 562–579. https://doi.org/10.1162/jocn.2006.18.4.562

Supèr, H., Romeo, A., & Keil, M. (2010). Feed-forward segmentation of figure-ground and assignment of border-ownership. PLoS ONE, 5(5), e10705. https://doi.org/10.1371/journal.pone.0010705

von der Heydt, R. (2015). Figure-ground organization and the emergence of proto-objects in the visual cortex. Frontiers in Psychology, 6(NOV), 1695. https://doi.org/10.3389/fpsyg.2015.01695

von der Heydt, R., Macuda, T., & Qiu, F. T. (2005). Border-ownership-dependent tilt aftereffect. Journal of the Optical Society of America A, 22(10), 2222. https://doi.org/10.1364/JOSAA.22.002222

Williford, J., & Heydt, R. (2013). Border-ownership coding. Scholarpedia, 8(10), 30040. https://doi.org/10.4249/scholarpedia.30040

Zhaoping, L. (2005). Border ownership from intracortical interactions in visual area V2. Neuron, 47(1), 143–153. https://doi.org/10.1016/j.neuron.2005.04.005

Zhou, H., Friedman, H. S., & von der Heydt, R. (2000). Coding of border ownership in monkey visual cortex. Journal of Neuroscience, 20(17), 6594. https://doi.org/10.1523/JNEUROSCI.2797-12.2013

Zipser, K., Lamme, V. A. F., & Schiller, P. H. (1996). Contextual Modulation in Primary Visual Cortex. The Journal of Neuroscience, 16(22), 7376–7389. https://doi.org/10.1523/JNEUROSCI.16-22-07376.1996

